# Serotonin Receptor Signaling Promotes Amoeboid Migration of Melanoma Cells Through cAMP/PKA Activation

**DOI:** 10.64898/2026.07.27.741069

**Authors:** Alleah G. Abrenica, Sylvia Kuang, Neelakshi Kar, Jeremy S. Logue

## Abstract

Melanoma cells frequently utilize bleb-driven amoeboid migration to navigate confined microenvironments during metastasis. Through a morphology-based drug repurposing screen, we identified the anti-migraine medication dihydroergotamine (DHE) as a potent inhibitor of this behavior. We show that DHE suppresses amoeboid motility by targeting serotonin receptors, primarily 5-HTR7. Trace levels of peripheral serotonin activates 5-HTR7, which in turn stimulates cAMP/PKA and ROCK signaling. This signaling cascade increases the cortical contractility necessary to sustain blebbing. Accordingly, DHE treatment or PKA inhibition significantly impairs confined migration without altering cell proliferation. Clinically, elevated levels of serotonin-degrading enzymes, such as MAOB, correlate with favorable patient prognoses and are reduced in metastatic lesions. Ultimately, these findings reveal an unanticipated role for peripheral serotonin in melanoma biomechanics and highlight serotonin receptors as highly actionable targets to limit metastatic dissemination.

**Significance Statement:** Metastasis is the leading cause of melanoma-related mortality, driven in part by the ability of tumor cells to adopt a highly contractile amoeboid mode of migration that enables movement through spatially confined tissues. This study uncovers a previously unrecognized mechanobiological role for peripheral serotonin in promoting this invasive behavior. We show that low concentrations of serotonin activate cAMP/PKA signaling to generate the actomyosin contractility required for amoeboid migration. Importantly, the FDA-approved migraine drug dihydroergotamine (DHE) blocks this pathway and significantly inhibits cancer cell motility. Supported by patient data linking reduced serotonin degradation with increased metastatic progression, these findings identify serotonin signaling as a novel driver of melanoma dissemination and provide a compelling rationale for repurposing serotonin-targeting therapeutics to prevent metastatic spread.

## Introduction

Cell migration is essential for development, wound healing, and immune surveillance; however, deregulated migration in cancer promotes metastasis. To fulfill targeted functions, cells may rely on distinct modes of movement. For instance, fibroblasts may adopt a mesenchymal mode of migration, characterized by the protrusion of an F-actin rich leading edge and the ligation of integrins to adhesion proteins [1]. Other populations, such as specific embryonic lineages, immune cells, and cancer cells, frequently utilize an amoeboid mode [2]. This type of migration is characterized by the formation of intracellular pressure-driven blebs and a lack of integrin-mediated adhesion [3]. Although unconstrained tissue environments abundant in adhesive ligands favor mesenchymal migration, narrow confinement and diminished adhesion prompt a shift toward amoeboid motility.

While cells are capable of navigating collagen matrices using small, dynamic blebs through mechanical “worrying,” rapid amoeboid migration relies on the formation of a single, large, stable bleb, termed a leader bleb [4]. When vertically confined down to ∼3 microns, cells may undergo a phenotypic transition to leader bleb-based migration (LBBM) [5]. This movement is powered by a rapid retrograde cortical actin flow, sustained by a concentration of myosin at the leader bleb neck [5–7]. The transmission of force to the microenvironment only requires friction; therefore, cells adopting fast amoeboid (leader bleb-based) migration are capable of moving through highly diverse environments [8]. Because this mode of migration requires elevated intracellular pressure, cells with profound actomyosin contractility are more likely to undergo this transition [6, 9]. For instance, intravital imaging reveals that cancer cells adopt an amoeboid phenotype at the invasive edges of tumors [10]. The dense fibrotic stroma surrounding tumors and the abundance of certain inflammatory factors (e.g., TGF-β) promote the shift to an amoeboid phenotype at invasive edges [11].

In the presence of adhesive ligands, cells may rapidly switch between phenotypes or adopt a hybrid state (i.e., displaying both mesenchymal and amoeboid features), as frequently observed in confining channels coated with VCAM-1 to simulate the microcapillary lumen [12]. In melanoma cells, the Piezo1/Ca(2+) pathway has been shown to sense confinement and trigger the amoeboid transition by activating inverted formin 2 (INF2) [12]. INF2 activation induces an “actin reset,” which promotes focal adhesion disassembly and the mesenchymal-to-amoeboid transition [12, 13]. The capacity to rapidly shift between mesenchymal and amoeboid phenotypes allows melanoma cells to traverse heterogeneous environments and drive metastasis, a migratory adaptability rooted in their neural crest origins.

Therefore, blocking the adoption of amoeboid features (i.e., blebs) may hold significant therapeutic value [14, 15]. Previously, we described a morphology-based drug repurposing screen in which we identified several drugs with anti-blebbing activity [16]. Among them, we identified dihydroergotamine (DHE), a prescription medication used to treat migraines. The tetracyclic ergoline core resembles the indole ring of serotonin (5-HT), making the serotonin receptors the primary targets of DHE. Here, we detail the novel pathway by which DHE inhibits the amoeboid (i.e., bleb-based) migration of melanoma cells, thus providing a new approach to limit metastatic dissemination.

## Results

To identify druggable pathways regulating blebbing, we previously performed a morphology-based drug repurposing screen [16]. While several drugs demonstrated anti-blebbing activity in A375 cells, here we focused on the prescription anti-migraine drug dihydroergotamine (DHE) because of its well-established safety profile (Fig. 1A). To confirm the results of our screen, we first subjected A375 cells to 2-dimensional (2D) or vertical confinement to induce fast amoeboid (leader bleb-based) migration [17]. In this assay, cells are confined by a PDMS ceiling coated with bovine serum albumin (1%) down to 3 µm, which is the optimal height at which cells undergo this phenotypic transition (Fig. 1B & Movie S1) [6]. Relative to DMSO (vehicle control), cells treated with DHE (10 µM) are far less likely to adopt a leader mobile (LM) phenotype (Fig. 1C-D). Consequently, DHE treated cells migrate significantly slower (Fig. 1E). Evaluating the kinematics of only motile cells reveals relatively modest changes in speed and directionality (Fig. 1E-F). Detailed morphological analyses show that DHE treatment significantly reduces both the largest bleb area and aspect ratio, while increasing roundness and circularity, which is consistent with a reduction in the number of leader mobile cells (Fig. 1D & S1A) [18]. We then used transmigration assays in which cells must migrate through 8 µm confining pores towards a fetal bovine serum (FBS; 10%) gradient. Relative to DMSO (vehicle control), DHE treatment led to a large reduction in transmigration (Fig. 1G). Similarly, DHE significantly inhibits the transmigration of another patient-derived melanoma cell line (WM983B) (Fig. 1H). Thus, DHE has significant activity against the confined migration of melanoma cells.

**Figure 1.**
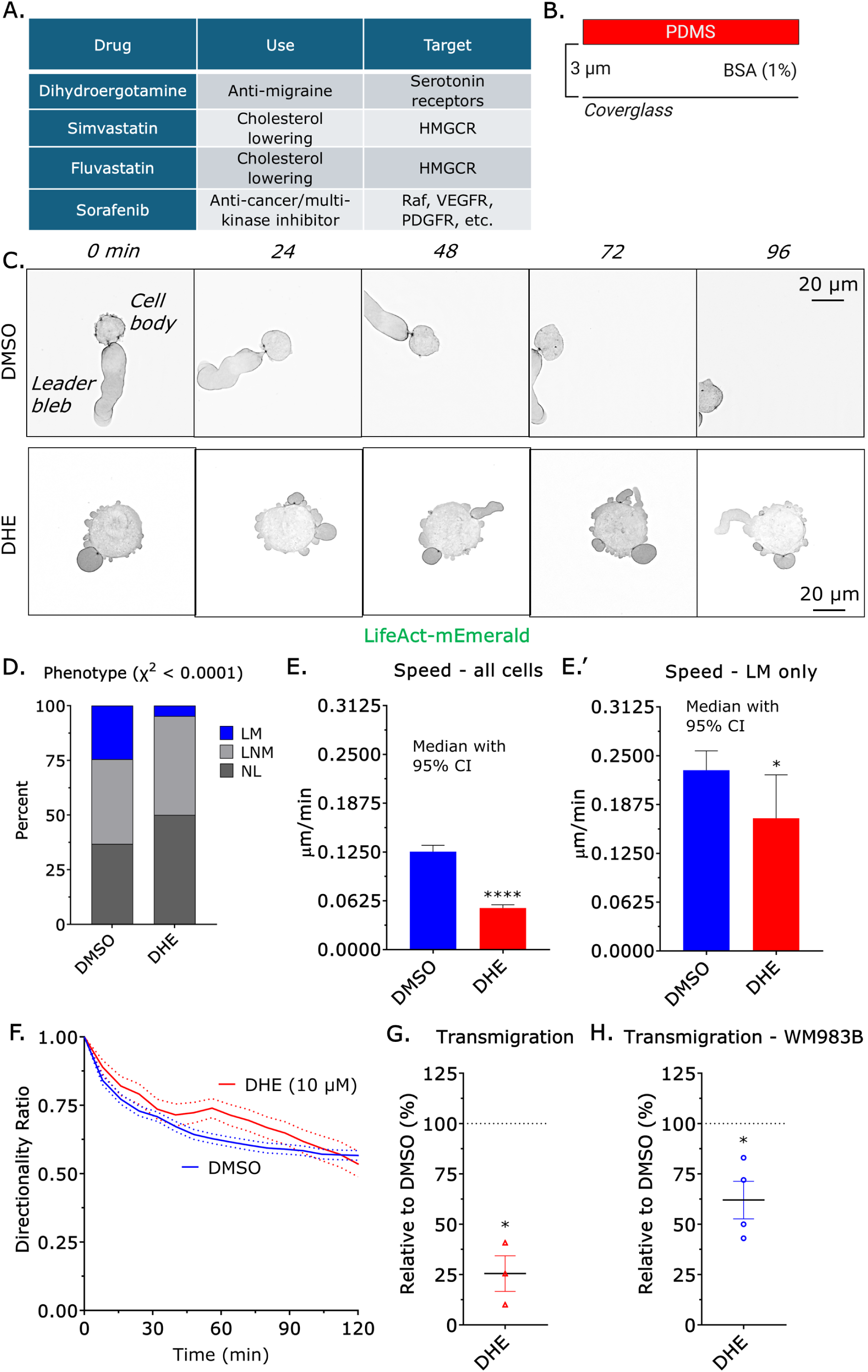
DHE inhibits leader bleb-based migration. **A.** Selected drugs, their clinical uses, and identified targets showing anti-blebbing activity in our morphological screen. **B.** Vertical confinement setup used to induce leader bleb-based migration. Cells are compressed beneath a BSA coated (1%) PDMS ceiling, maintained at a defined height using 3 µm beads. **C.** Representative montages of A375 cells treated with DMSO or DHE. To visualize F-actin, cells were transiently transfected with LifeAct-mEmerald. Control cells adopt a leader mobile (LM) phenotype, whereas DHE-treated cells predominantly display a non-leader (NL) morphology. **D.** Distribution of LM, leader non-mobile (LNM), and NL phenotypes in cells treated with DMSO (*n* = 754) or DHE (*n* = 681). Statistical significance was determined using a χ^2^ test. **E.** Instantaneous speeds of all tracked cells treated with DMSO (*n* = 754) or DHE (*n* = 681). All phenotypic subsets were included. Statistical significance was evaluated by a Mann-Whitney test. **E’.** Instantaneous speeds isolated to the LM phenotype for DMSO (*n* = 204) and DHE (*n* = 38) conditions. Statistical significance was evaluated by a Mann-Whitney test. **F.** Directionality ratio (d/D) over time for both treatment groups. Data are shown as mean +/- SEM. **G - H.** DHE diminishes the transmigration of A375 (G) and WM983B (H) cells through 8 µm pores toward an FBS (10%) gradient, relative to DMSO controls. Data are shown as mean +/- SEM. Statistical significance was determined by a one-sample *t*-test against a hypothetical value of 100%. Significance levels: * - p ≤ 0.05, ** - p ≤ 0.01, *** - p ≤ 0.001, and **** - p ≤ 0.0001.

Ergoline compounds broadly display low nanomolar affinity to the serotonin (5-HT) family of receptors [19]. Because melanocytes emerge from the neural crest, their transcriptome is inherently programmed with signature neuronal pathways. Therefore, by reverse transcriptase quantitative PCR (qPCR), we determined the relative levels of all known serotonin receptors in A375 cells. Relative to GAPDH, the predominant transcript in these cells encodes 5-HTR7, a Gs and Gα12/13 coupled receptor robustly inhibited by DHE at low nanomolar concentrations (Fig. 2A) [20, 21]. Notably, 5-HTR7 was also the most abundant family member in WM983B cells (Fig. S2A). As cells in tissues traverse a range of confined environments, we determined if DHE inhibits migration through microchannels (measuring 8 µm by 8 µm by 100 µm) coated with VCAM-1 (1 µg/mL) to simulate the microcapillary lumen (Fig. 2B-C & Movie S2). Under these conditions, A375 cells adopt an amoeboid (i.e., blebbing), mesenchymal, or hybrid phenotype. Relative to DMSO (vehicle control), DHE treatment led to a significant reduction in all migration modes (Fig. 2D). Upon analyzing only motile cells, we also observed significant decreases in both speed and directionality (Fig. 2E-F). While DHE treatment inhibited migration, it had little effect on the rate of proliferation in A375 cells (Fig. 2G). To confirm that 5-HTR7 is the primary target of DHE, we used RNAi. Relative to non-targeting controls, RNAi of 5-HTR7 led to a significant reduction in the number of motile cells (Fig. 2H-I). Strikingly, the largest reduction was in the number of amoeboid cells (Fig. 2H-I). Following up on this result, we confined cells in channels coated with bovine serum albumin (BSA; 1%), which by blocking adhesion promotes the amoeboid phenotype. Using this approach, we observed nearly a twofold reduction in motile amoeboid cells (Fig. 2J). Additionally, RNAi of 5-HTR7 inhibited transmigration through confining pores (Fig. 3A). Collectively, these data indicate that 5-HTR7 is the primary target of DHE in A375 and WM983B cells, which specifically promotes the amoeboid phenotype in confining channels.

**Figure 2.**
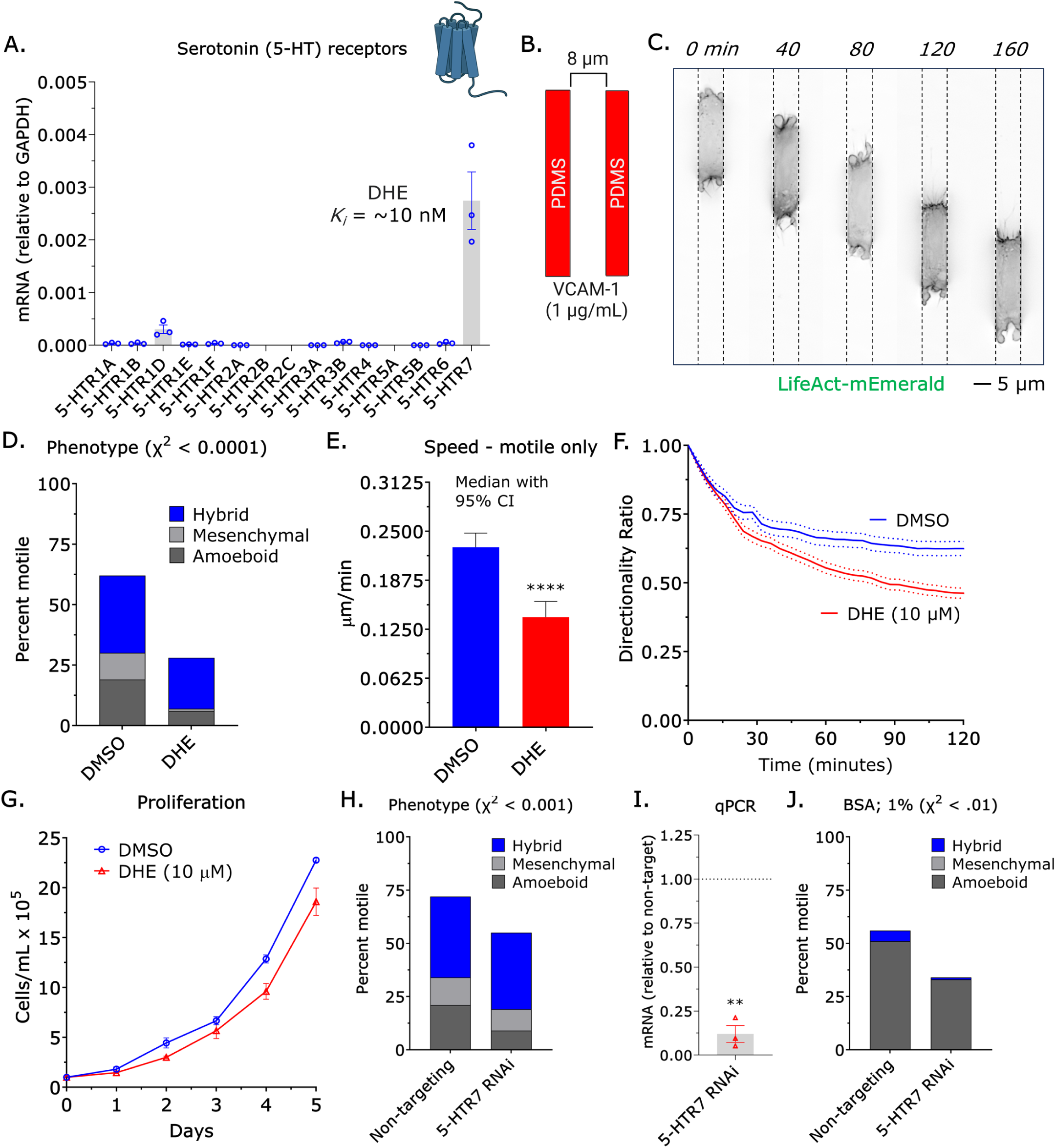
DHE inhibits the migration of melanoma cells through confining channels. **A.** The 5-HTR7 transcript is abundant in A375 cells, as quantified by reverse transcriptase qPCR. Data represent the mean +/- SEM. **B.** Microchannel setup for inducing amoeboid migration. Cells spontaneously migrate through channels (8 by 8 by 100 µm) coated with VCAM-1 (1 µg/mL). C. Representative montage illustrating the hybrid phenotype, wherein cells demonstrate both amoeboid (*i.e.*, blebs) and mesenchymal features. Cells were transiently transfected with mEmerald-LifeAct to visualize F-actin. **D.** Distribution of hybrid, mesenchymal, and amoeboid phenotypes following treatment with DMSO (*n* = 329) or DHE (*n* = 212). Statistical significance was evaluated using a χ^2^ test. **E.** Instantaneous speeds of motile cells for DMSO (*n* = 204) and DHE (*n* = 59) conditions. Statistical significance was determined by a Mann-Whitney test. **F.** Directionality ratio (d/D) over time for both treatment groups. Data represent the mean +/- SEM. **G.** Cellular proliferation over time for both treatment groups. Data represent the mean +/- SEM. **H.** Distribution of hybrid, mesenchymal, and amoeboid phenotypes in cells treated with a non-targeting (*n* = 242) or 5-HTR7 siRNA (*n* = 141). Statistical significance was evaluated using a χ^2^ test. **I.** Relative to non-targeting controls, 5-HTR7 transcript levels are significantly diminished by RNAi, as measured by reverse transcriptase qPCR. Data represent the mean +/- SEM. Statistical significance was determined by a one-sample *t*-test against a hypothetical value of 1. **J.** Distribution of hybrid, mesenchymal, and amoeboid phenotypes in cells treated with a non-targeting (*n* = 81) or 5-HTR7 siRNA (*n* = 81). To selectively promote amoeboid migration, channels were coated with BSA (1%). Statistical significance was determined using a χ^2^ test. Significance levels: * p ≤ 0.05, p ≤ 0.01, *** p ≤ 0.001, and **** p ≤ 0.0001.

**Figure 3.**
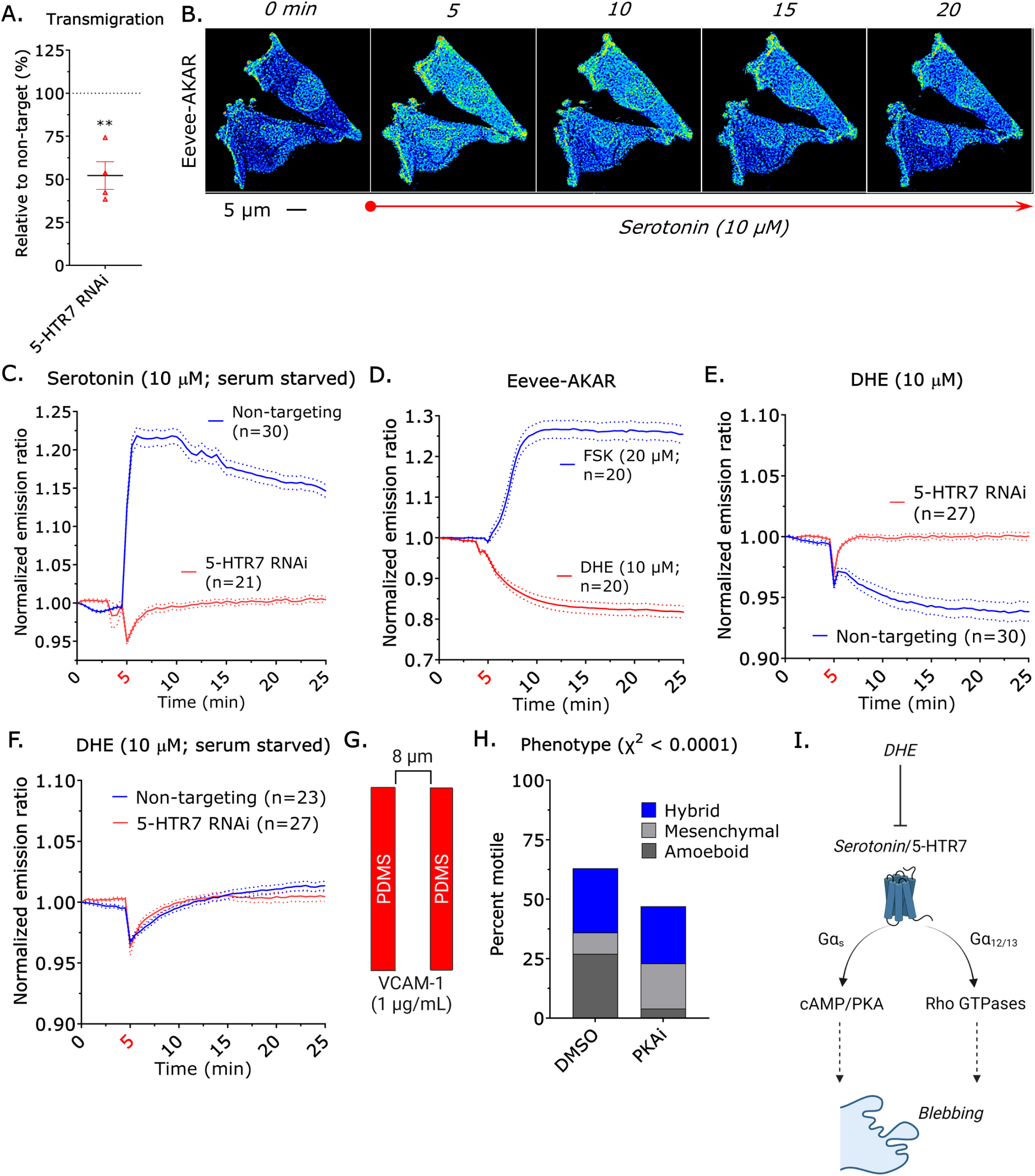
DHE blunts cAMP/PKA signaling downstream of 5-HTR7. **A.** RNAi of 5-HTR7 attenuates the transmigration of A375 cells through 8 µm pores toward an FBS (10%) gradient, relative to non-targeting controls. Data represent the mean +/- SEM. Statistical significance was determined by a one-sample *t*-test against a hypothetical value of 100%. **B.** Representative montage of Eevee-AKAR. A warmer pseudo color indicates heightened PKA activity, as measured by FRET. Serotonin was added to serum-starved cells at 5 min. **C.** RNAi of 5-HTR7 prevents the activation of PKA by serotonin, relative to non-targeting controls. Serotonin was added to serum-starved cells at 5 min. Data represent the mean +/- SEM. **D.** The addition of forskolin (FSK) induces cAMP/PKA signaling, whereas the addition of DHE reduces it. Drugs were added to cells in complete media at 5 min. Data represent the mean +/- SEM. **E.** RNAi of 5-HTR7 diminishes the response to DHE, relative to non-targeting controls. DHE was added to cells in complete media at 5 min. **F.** The inhibition of PKA activity downstream of 5-HTR7 requires serum serotonin. DHE was added to serum-starved cells at 5 min. Data represent the mean +/- SEM. **G.** Microchannel setup for inducing amoeboid migration. Cells spontaneously migrate through channels (8 by 8 by 100 µm) coated with VCAM-1 (1 µg/mL). **H.** Distribution of hybrid, mesenchymal, and amoeboid phenotypes following treatment with DMSO (*n* = 241) or a PKA inhibitor (KT5720; *n* = 210). Statistical significance was evaluated using a χ^2^ test. **I.** Illustration of the two signaling arms activated downstream of 5-HTR7 by serotonin, which may regulate blebbing. Significance levels: * - p ≤ 0.05, ** - p ≤ 0.01, *** - p ≤ 0.001, and **** - p ≤ 0.0001.

Having established that 5-HTR7 promotes migration in 2D and 3D (e.g., microchannels) confined environments, we sought to determine whether serotonin triggers the activation of cAMP/PKA signaling in A375 cells. To answer this, we used the Eevee-A kinase activity reporter (AKAR) [22]. In serum-starved A375 cells, we observed a very rapid increase in the normalized emission ratio (i.e., FRET) after the addition of 10 µM serotonin (Fig. 3B-C & Movie S3). However, RNAi blocked this effect, confirming that the activation of cAMP/PKA signaling by serotonin requires 5-HTR7 in A375 cells (Fig. 3B-C). Using 20 µM forskolin (FSK) to directly activate adenylyl cyclase, we confirmed that Eevee-AKAR responds to a rise in cAMP/PKA activity, which was similar in magnitude to serotonin (Fig. 3C-D). Serotonin activity could be blocked by a PKA inhibitor (KT5720), as measured by Eevee-AKAR (Fig. S3A). In contrast, DHE treatment rapidly decreased the emission ratio (Fig. 3D). However, DHE had no effect on cells lacking 5-HTR7, which is consistent with the idea that this receptor is the primary target of DHE (Fig. 3E). Additionally, DHE had no effect on serum-starved cells (Fig. 3F). While serotonin can drive cellular behavior through autocrine loops, this does not appear to happen in A375 cells. To evaluate potential autocrine production, we quantified the transcripts of TPH1 and TPH2, the fundamental enzymes for serotonin synthesis. By reverse transcriptase qPCR, A375 cells show virtually no transcripts for either enzyme (Fig. S4A) [23]. Corroborating this finding, conditioned media could not initiate cAMP/PKA signaling in serum-starved cells (Fig. S4B). Thus, rather than utilizing an autocrine loop, these melanoma cells are stimulated by the ambient serotonin inherently present in serum (Fig. S4B) [24]. Crucially, we found that the addition of a PKA inhibitor inhibited migration in 2D and 3D (e.g., microchannels) confined environments (Fig. 3G-H & S3B-C). Interestingly, in confined channels coated with VCAM-1 (1 µg/mL), the number of cells adopting an amoeboid phenotype was significantly reduced, whereas the number of mesenchymal cells was significantly increased after inhibiting PKA (Fig. 3G-H). While these data demonstrate that cAMP/PKA signaling promotes amoeboid (i.e., bleb-based) migration, 5-HTR7 also activates Rho by way of Gα12/13 (Fig. 3I).

Serotonin may promote the amoeboid migration of melanoma cells by activating both cAMP/PKA and Rho, as 5-HTR7 is coupled to both G_s_ and Gα12/13 [20, 21]. This idea is supported by reverse transcriptase qPCR, which shows significant amounts of both Gα12 and Gα13 transcript in A375 cells (Fig. 4A). In serum-starved cells, serotonin was found to rapidly activate ROCK, as measured by the emission ratio of Eevee-ROCK (Fig. 4B-C & Movie S4) [25]. Strikingly, this effect was highly similar in magnitude to stimulation with 10% serum (Fig. 4C). However, RNAi of 5-HTR7 blocked the effect of serotonin, further confirming that this receptor is the primary target of serotonin (Fig. 4D). Given that serotonin was found to potently activate each signaling pathway, we determined if serotonin alone (i.e., in the absence of serum) is sufficient to drive migration through confining pores. Indeed, we found that serotonin was sufficient to drive transmigration, which could be blocked by the addition of DHE (Fig. 4E). Remarkably, this effect was similar in magnitude to serum (Fig. 4E). However, serum stimulation was still able to activate ROCK after RNAi of 5-HTR7, which demonstrates that other serum factors can activate ROCK in the absence of this receptor (Fig. 4F). Consistent with this data, ROCK activity was modestly reduced by DHE in cells in complete media, whereas the addition of a ROCK inhibitor (Y27632) led to a large reduction (Fig. 4G). In A375 cells, the activity of ROCK was unaffected by the addition of a PKA inhibitor; therefore, ROCK activity is not sustained by PKA (Fig. 4G). While these data demonstrate that serotonin is sufficient to activate ROCK, they also show that it is not necessary, as other serum factors can activate ROCK in the absence of 5-HTR7. As cortical contractility drives amoeboid (i.e., bleb-based) migration, we used a previously described assay to directly measure the stiffness of cells. We compressed spherical cells between polyacrylamide gels of known stiffness (1 kPa) (Fig. 4H). For each cell, we determined by microscopy its height (*h*) and diameter (*d*), which were used to calculate a ‘stiffness’ (*h*/*d*). Relative to DMSO (vehicle control), DHE treated cells displayed a significant reduction in cell stiffness (Fig. 4I). This reduction was similar in magnitude to treatment with a PKA inhibitor (KT5720), whereas treatment with a ROCK inhibitor (Y27632) had the largest effect (Fig. 4I). In contrast, serotonin treatment led to a significant increase in cell stiffness (Fig. 4I). In agreement with these data, RNAi of 5-HTR7 also led to a significant decrease in cell stiffness (Fig. 4J). Although PKA has many substrates in cells, RNAi of a well-known substrate, VASP, led to a significant reduction in stiffness (Fig. 4K) [26]. Therefore, PKA signaling is sufficient to drive the cortical contractility required for amoeboid migration.

**Figure 4.**
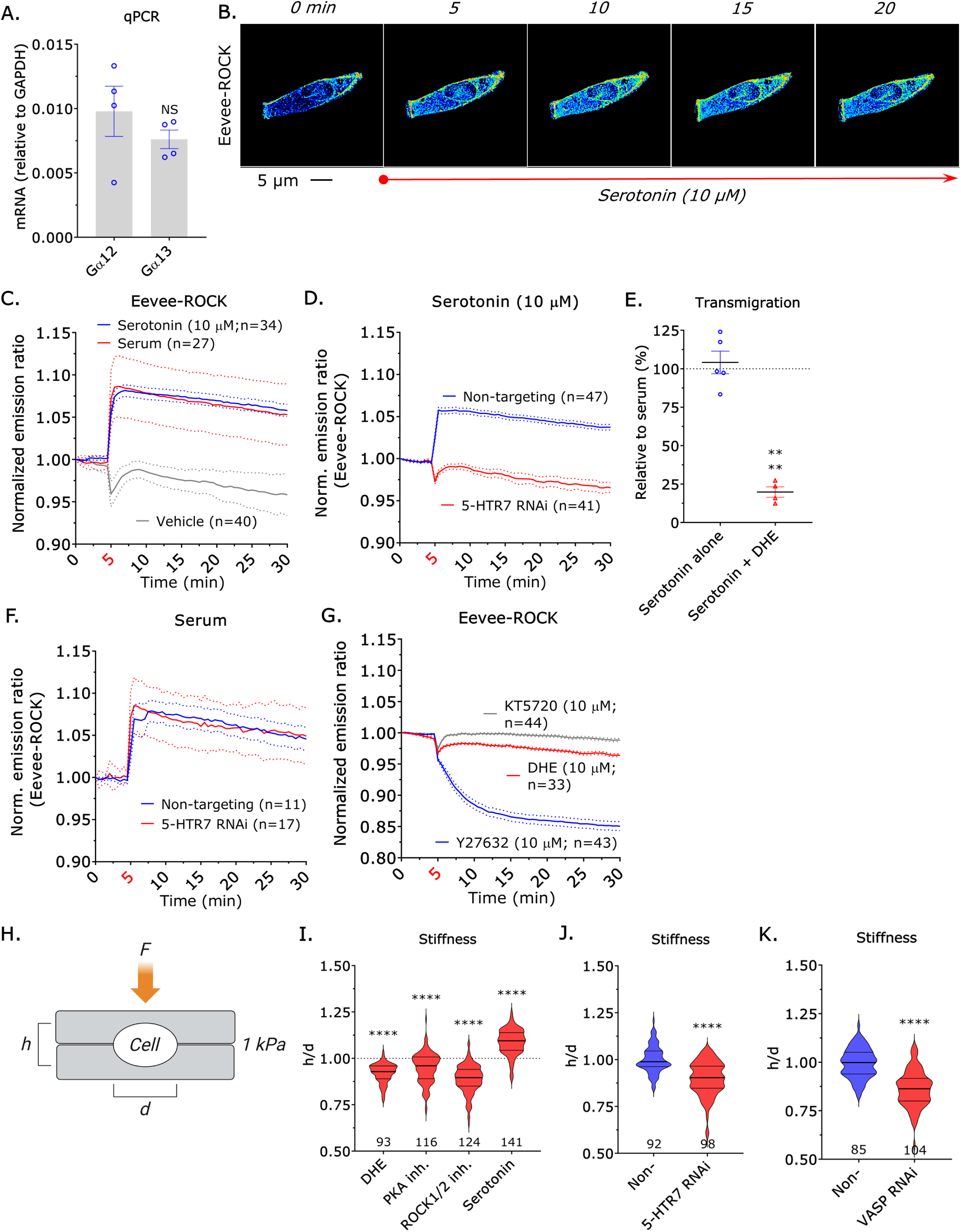
5-HTR7 mediated signaling drives cortical contractility. **A.** High levels of Gα12 and Gα13 transcripts are present in A375 cells, quantified by reverse transcriptase qPCR. Data represent the mean +/- SEM. Statistical significance was evaluated by a Welch’s *t*-test. **B.** Representative montage of the Eevee-ROCK biosensor. A warmer pseudocolor indicates elevated ROCK activity, evaluated by FRET. Serotonin was introduced to serum-starved cells at 5 min. **C.** Introduction of serotonin or serum (10%) yields a comparable induction of ROCK activity, relative to the vehicle control. Both were administered to serum-starved cells at 5 min. Data represent the mean +/- SEM. **D.** RNAi targeting 5-HTR7 prevents ROCK activation by serotonin, relative to non-targeting controls. Serotonin was administered to serum-starved cells at 5 min. Data represent the mean +/- SEM. **E.** Serotonin stimulates cell transmigration through 8 µm pores, relative to serum (10%) controls. Co-administration of serotonin and DHE abolishes this stimulatory effect, relative to serum (10%) controls. Data represent the mean +/- SEM. Statistical significance was determined by a Welch’s *t*-test. **F.** RNAi targeting 5-HTR7 fails to inhibit ROCK activation by serum (10%), relative to non-targeting controls. Serum (10%) was administered to serum-starved cells at 5 min. Data represent the mean +/- SEM. **G.** Compared to Y27632, adding DHE at 5 min to cells in complete media induces a moderate decline in ROCK activity. Adding a PKA inhibitor (KT5720) produced no measurable effect on ROCK activity. Data represent the mean +/- SEM. **H.** Gel sandwich assay used to deform cells between 1 kPa polyacrylamide gels. Stiffness is calculated as the ratio of cell height (*h*) to diameter (*d*). **I.** Cellular stiffness following a 1 hr treatment with DHE (10 µM), a PKA inhibitor (KT5720; 10 µM), a ROCK1/2 inhibitor (Y27632; 10 µM), or serotonin (10 µM), relative to vehicle controls. The values below each plot indicate the total number of cells analyzed. The median and interquartile ranges are provided. Statistical significance was determined by a one-sample *t*-test against a hypothetical value of 1. **J. – K.** Cellular stiffness following RNAi targeting 5-HTR7 (J) or VASP (K), relative to non-targeting controls. The median and interquartile ranges are provided. Statistical significance was evaluated by a Welch’s *t*-test. Significance levels: * - p ≤ 0.05, ** - p ≤ 0.01, *** - p ≤ 0.001, and **** - p ≤ 0.0001.

Having established that 5-HTR7 signaling drives amoeboid (i.e., bleb-based) migration in A375 and WM983B, we wondered if other melanoma “workhorse” cell lines might be sensitive to DHE. The Yale University Mouse Melanoma (YUMM) 1.7 and 3.3 cells, due to their defined genetic background, are heavily used to generate syngeneic tumors in C57BL/6 mice [27]. Interestingly, we could not detect 5-HTR7 transcripts in YUMM1.7 or YUMM3.3 cells, as measured by reverse transcriptase PCR (Fig. 5A-B). However, in YUMM3.3 cells we could detect significant levels of 5-HTR1B, whereas YUMM1.7 cells display minimal 5-HTR1B transcript (Fig. 5A-B). While 5-HTR7 is detected at much higher levels, 5-HTR1B transcript is detected in patient tumors (Fig. 5C & S5A). However, 5-HTR1B is Gi coupled (Fig. 5C) [28]. Crucially, DHE is known to bind with sub-nanomolar affinity to 5-HTR1B. As opposed to the competitive antagonist activity against 5-HTR7, DHE is a 5-HTR1B agonist [29]. Thus, DHE similarly inhibits PKA signaling in cells with 5-HTR1B. In agreement with this concept, DHE treatment led to a significant reduction in the number of cells adopting a leader mobile (LM) phenotype (Fig. 5D). Similarly, transmigration through confining pores was significantly reduced by DHE, whereas the proliferation of YUMM3.3 cells was unaffected (Fig. 5E-F). Detailed morphometric analyses revealed a decrease in both the largest bleb area and aspect ratio, alongside an increase in both roundness and circularity in cells treated with DHE, which is consistent with a reduction in the number of LM cells (Fig. 5D & S6A). Remarkably, due to the “promiscuous” nature of DHE, it is effective against amoeboid migration in cell models with varying serotonin receptor complements.

**Figure 5.**
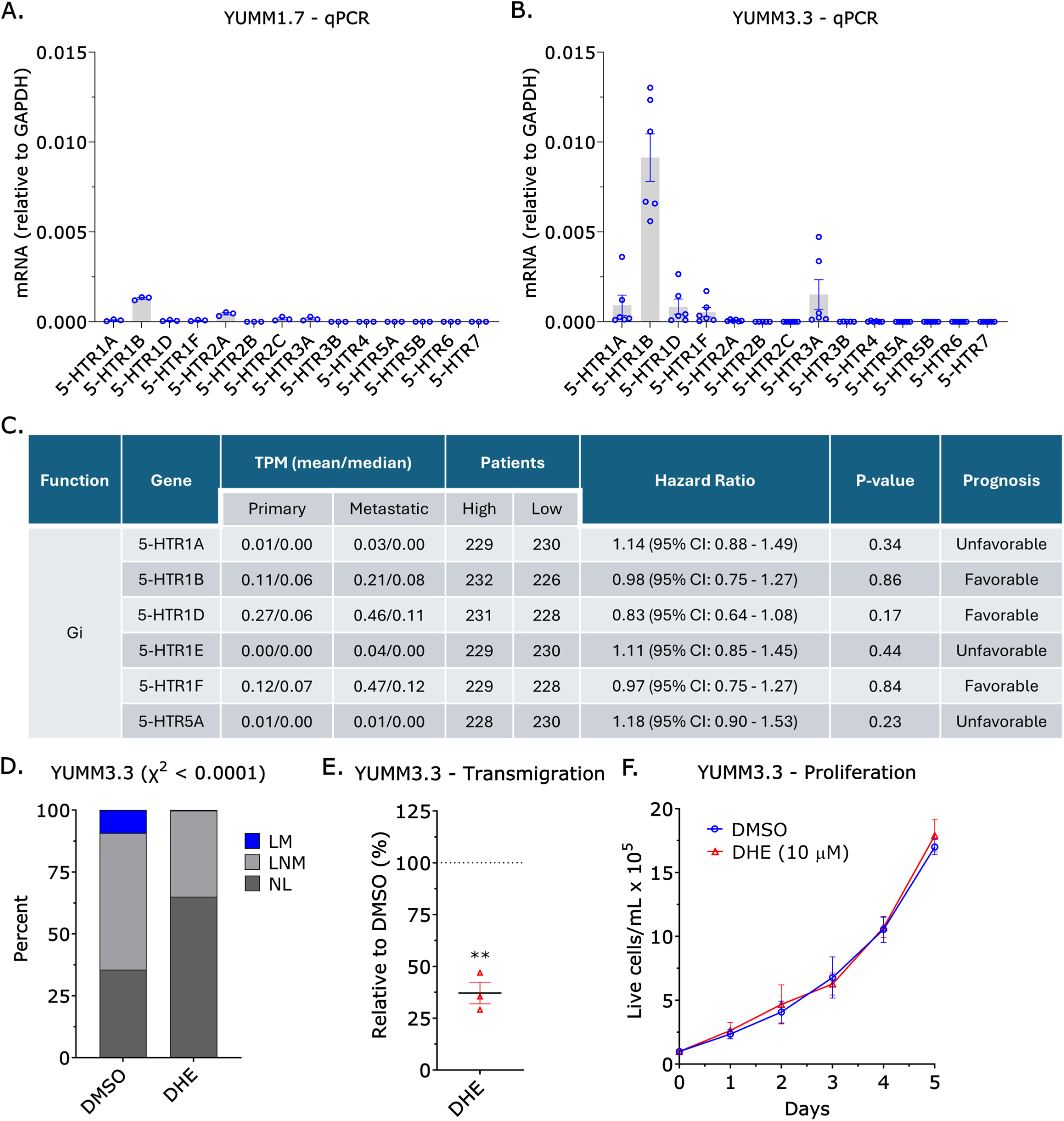
Multiple melanoma cell lines remain sensitive to DHE despite harboring distinct serotonin receptor profiles. **A.** Serotonin receptor transcript levels are minimal in YUMM1.7 cells, quantified by reverse transcriptase qPCR. Data represent the mean +/- SEM. **B.** Abundant 5-HTR1B transcript levels are present in YUMM3.3 cells, quantified by reverse transcriptase qPCR. Data represent the mean +/- SEM. **C.** Tabulation of Gi coupled serotonin receptors detailing transcripts per million (TPM) values within primary and metastatic tumors. Additional details include the number of patients stratified by serotonin receptor transcript levels, hazard ratios, p-values, and overall prognoses. Transcriptomic data for cutaneous melanoma were sourced from The Cancer Genome Atlas through the Genomic Data Commons. **D.** Distribution of leader mobile (LM), leader non-mobile (LNM), and no leader (NL) phenotypes within YUMM3.3 cells treated with DMSO (*n* = 164) or DHE (*n* = 238). Statistical significance was evaluated by a χ^2^ test. **E.** DHE attenuates the transmigration of YUMM3.3 cells through 8 µm pores toward an FBS (10%) gradient, relative to DMSO controls. Data represent the mean +/- SEM. Statistical significance was determined by a one-sample *t*-test against a hypothetical value of 100%. **F.** Cellular proliferation evaluated over time for both treatment groups. Data represent the mean +/- SEM. Significance levels: * - p ≤ 0.05, ** - p ≤ 0.01, *** - p ≤ 0.001, and **** - p ≤ 0.0001.

Having established that 5-HTR7 signaling promotes amoeboid (i.e., bleb-based) migration in patient-derived melanoma cells, we set out to determine if the levels of any serotonin signaling-associated factors correlate with disease severity. To this end, we utilized publicly available transcriptomic data (e.g., The Cancer Genome Atlas) for cutaneous melanoma. While a high level of 5-HTR7 was not associated with an unfavorable prognosis, high levels of MAOB and ALDH2 were associated with a favorable prognosis (Fig. 6A & S6A). Each of these enzymes participates in the degradation of serotonin; therefore, high levels of MAOB or ALDH2 within the tumor microenvironment (TME) are likely to reduce serotonin levels. Relative to the primary, the mean and median transcripts per million (TPM) for MAOB are reduced in metastatic tumors (Fig. 6A). To validate this observation at the protein level, we stained melanoma tissue arrays for MAOB. Using quantitative immunofluorescence imaging, we observed a significant decrease in the level of MAOB in metastatic tumors (Fig. 6B-C). Thus, a reduction in the level of a key serotonin-degrading enzyme within the TME is correlated with metastatic dissemination.

**Figure 6.**
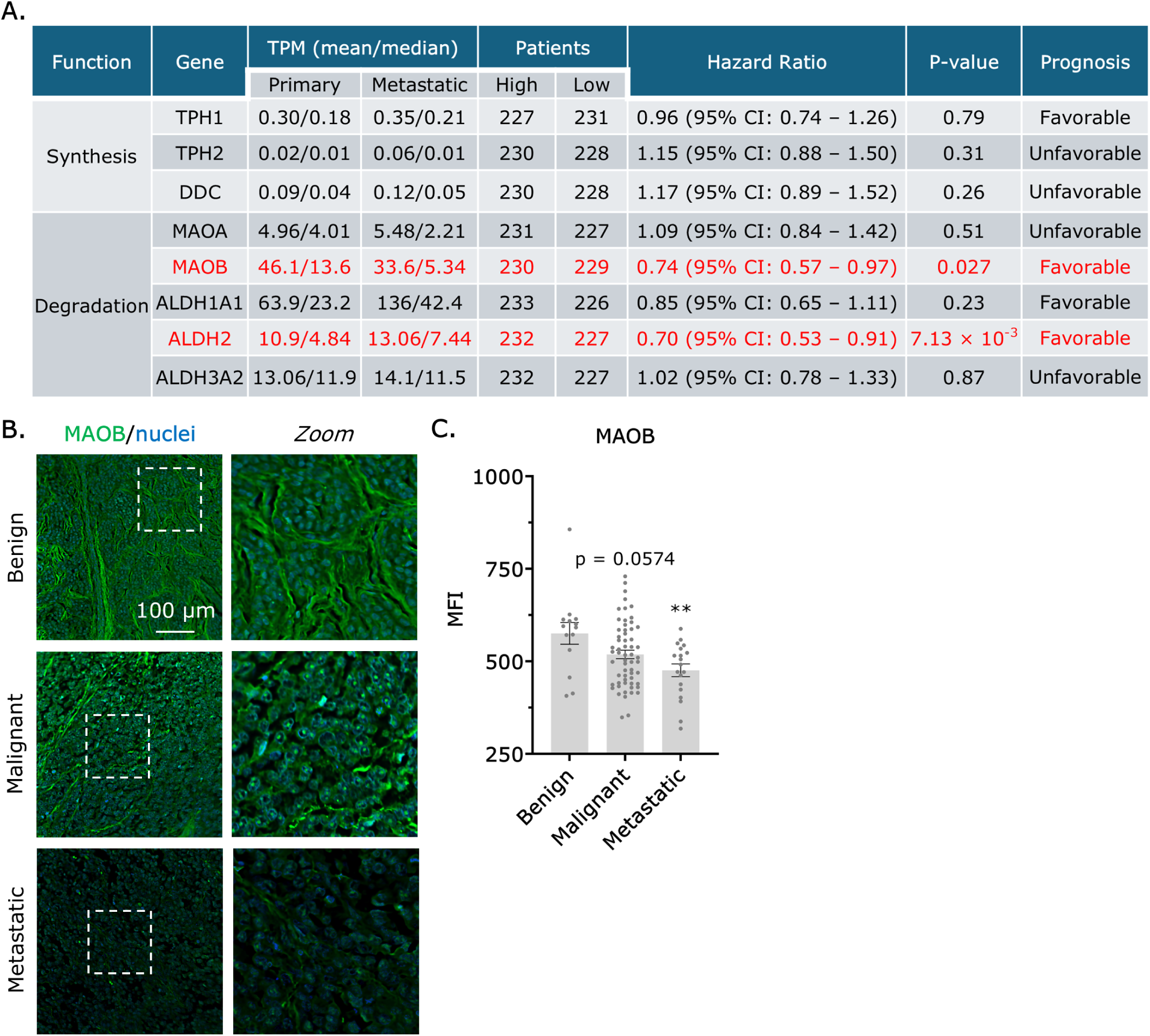
The abundance of enzymes governing serotonin degradation correlates with prognosis in melanoma patients. **A.** Tabulation of enzyme transcript levels governing serotonin synthesis and degradation within primary and metastatic tumors. Additional details include the number of patients stratified by enzyme transcript levels, hazard ratios, p-values, and overall prognoses. Transcriptomic data for cutaneous melanoma were sourced from The Cancer Genome Atlas through the Genomic Data Commons. Enzymes with significant hazard ratios are highlighted in red. **B.** Immunofluorescence staining of benign, malignant, and metastatic melanoma tissue array samples for MAOB. Samples were counterstained for nuclei. **C.** Mean fluorescence intensities (MFI) for MAOB (green fluorescence) in benign (*n* = 14), malignant (*n* = 19), and metastatic melanoma (*n* = 60) tissue array samples. Data represent the mean +/- SEM. Statistical significance was determined by a one-way ANOVA and a Dunnett’s multiple comparison test. Significance levels: * - p ≤ 0.05, ** - p ≤ 0.01, *** - p ≤ 0.001, and **** - p ≤ 0.0001.

## Discussion

While serotonin (5-HT) is often viewed through the lens of central neurotransmission, ∼95% of serotonin is synthesized, utilized, and degraded within peripheral compartments [30]. As neural crest derivatives, melanocytes possess the receptors that allow peripheral serotonin to act as a powerful morphogenetic regulator. In various cancers, serotonin signaling is known to activate canonical pro-survival and mitogenic pathways [31]. Accordingly, it was revealing that a drug containing a serotonin-like indole ring effectively suppressed membrane blebbing in melanoma cells. Thus, the role of serotonin signaling in melanoma may go beyond traditional survival mechanisms. Our findings uncover a new function for these receptors in driving amoeboid (i.e., bleb-based) migration.

Using a morphology based repurposing screen, we identified a number of FDA-approved drugs with activity against blebs in melanoma cells. Previously, we reported that statins inhibit blebbing through the reduction of plasma membrane cholesterol, which is required to transmit force to Piezo1. We demonstrated that melanoma cells require this cation specific channel for sensing increasing levels of confinement [12, 16]. Here, we have focused on an entirely new 5-HTR7 mediated pathway, identified as the primary target of DHE in two patient-derived melanoma cell lines. Remarkably, DHE demonstrated activity against amoeboid (i.e., bleb-based) migration in 2D and 3D (e.g., microchannels) environments. This activity was not an indirect effect of reduced cell health, as cell proliferation was unaffected by DHE. Therefore, DHE specifically inhibits cell migration.

Having established that our patient-derived melanoma cell lines have abundant levels of 5-HTR7, we measured downstream signaling using biosensors for their high temporal resolution. Using Eevee-AKAR, serotonin rapidly and robustly activated cAMP/PKA signaling in melanoma cells [22]. Crucially, adding DHE led to a substantial decrease in PKA activity, an effect dependent on 5-HTR7. This reduction was not observed in starved cells, which indicates that the trace amount of serotonin found in serum is sufficient to activate this receptor [24]. In agreement with these data, a highly specific PKA inhibitor potently reduced the number of cells adopting an amoeboid phenotype in confining channels. Interestingly, inhibiting PKA also led to a significant increase in the number of mesenchymal cells. Thus, a high level of PKA activity within these cells drives a shift toward an amoeboid phenotype.

As 5-HTR7 is also coupled to Gα12/13, we evaluated whether serotonin stimulates Rho GTPase signaling [20, 21]. Using a specific biosensor, we found that serotonin could activate ROCK. Remarkably, the activation was similar in magnitude to serum, and the effect was dependent on 5-HTR7. Consistent with these data, the transmigration of A375 cells through confining pores was stimulated by serotonin alone (i.e., in the absence of serum factors). However, in cells depleted of 5-HTR7 by RNAi, serum stimulation was still capable of activating ROCK. Thus, serotonin is sufficient but not necessary for ROCK activation. In agreement with this concept, DHE only moderately reduced ROCK activity in cells maintained in complete media. Despite this, we sought to determine if 5-HTR7 signaling could upregulate cortical contractility, which is a powerful driver of amoeboid migration. Using a previously established gel sandwich approach, DHE treated cells were found to be more deformable, whereas serotonin treated cells were stiffer [6]. Crucially, treatment with a specific PKA inhibitor resulted in a similar reduction in stiffness. Moreover, RNAi of VASP, which is a well-known PKA substrate, similarly reduced cell stiffness [26]. Thus, cAMP/PKA signaling is sufficient to maintain cortical contractility in these cells. Although PKA has many substrates, remodeling of the cortical actin cytoskeleton by phosphorylated VASP may underlie this effect [32].

Although abundant in A375 and WM983B cells, we could not detect significant levels of 5-HTR7 transcript in YUMM1.7 or YUMM3.3 cells. In YUMM3.3 cells, we found a substantial amount of 5-HTR1B transcript. Remarkably, treating YUMM3.3 cells with DHE reduced the number of cells adopting a leader mobile phenotype. A detailed morphometric analysis, which showed reductions in both the largest bleb area and aspect ratio, confirmed this result. Due to its “promiscuous” nature, DHE acts as a 5-HTR1B agonist [29]. Thus, the downstream effect of DHE on PKA activity is unchanged, as 5-HTR1B is a Gi coupled receptor [28]. In addition to highlighting the unique pharmacology of DHE, these results underscore the universal importance of cAMP/PKA signaling in promoting amoeboid (i.e., bleb-based) migration.

Using publicly available transcriptomic data, we found that high levels of two serotonin-degrading enzymes correlate with a favorable prognosis in melanoma patients. Relative to primary tumors, we confirmed that MAOB protein is reduced in metastatic lesions. Whether invading cells home to sites abundant in serotonin, whether high serotonin levels enhance cell fitness through metabolic reprogramming, or whether cells remodel the surrounding stroma to downregulate the transcription of this gene remain as unanswered questions to be addressed in future studies. Collectively, our data demonstrate that 5-HTR7 signaling promotes the amoeboid migration of melanoma cells, thereby revealing a previously unknown role for peripheral serotonin in metastasis.

## Resource Availability

### Lead contact

Requests for further information should be directed to and will be fulfilled by the lead contact, Jeremy S. Logue.

### Materials availability

No unique/stable reagents were generated in this study.

### Data and code availability

Data and code used in the study are included in the manuscript and/or supporting information.

## Acknowledgments

We thank the Cady Lab (SUNY Polytechnic Institute, Albany, NY) for fabricating silicon wafer molds. This work was supported by grants from the Melanoma Research Alliance (MRA; award no. 688232) (DOI: https://doi.org/10.48050/pc.gr.91570), the American Cancer Society (ACS; award no. RSG-20-019-01 - CCG), and the National Institutes of Health (NIH; award no. R35GM146588) to J.S.L.

## Author Contributions

A.A.: Investigation, Writing – Original Draft, S.K.: Investigation, N.K.: Investigation, J.S.L.: Conceptualization, Supervision, Project administration, Visualization, Writing - Original Draft, Funding acquisition

## Declaration of Interests

The authors declare no competing interests.

## MATERIALS AND METHODS

### Cell lines

A375-M2 (CRL-3223), YUMM1.7 (CRL-3362), and YUMM3.3 (CRL-3365) were obtained from the American Type Culture Collection (Manassas, VA). WM983B cells (WM983B-01-0001) were obtained from Rockland Immunochemicals (Pottstown, PA). Cells were cultured in high-glucose DMEM supplemented with fetal bovine serum (FBS; 10%) (Sigma Aldrich; 12106C), L-glutamine (Thermo Fisher; 35050061), pyruvate, HEPES, and Antibiotic-Antimycotic (Thermo Fisher; 15240096).

### Plasmids

LifeAct-mEmerald (plasmid no. 54148) was obtained from Addgene (Watertown, MA). Eevee-AKAR and Eevee-ROCK was kindly provided by Dr. Michiyuki Matsuda (Kyoto University). 1 µg of plasmid was used to transfect 750,000 cells in each well of a 6-well plate using Lipofectamine 2000 (5 µL; Thermo Fisher) in OptiMEM (400 µL; Thermo Fisher). After 20 min at room temperature, plasmid in Lipofectamine 2000/OptiMEM was incubated with cells in complete media (2 mL) overnight.

### Chemical treatments

Dihydroergotamine mesylate (DHE; 475), Serotonin hydrochloride (5-HT; 3547), PKA activator (forskolin; 1099), PKA inhibitor (KT5720; 1288), and ROCK inhibitor (Y27632; 1254) were purchased from Tocris Bioscience (Bristol, UK). Prior to visualizing confined migration, devices were pre-incubated with compounds in complete media for at least 1 hr before loading cells. For serum-starved conditions, cells were incubated in serum-free media for ∼2 hr.

### RNA interference

Non-targeting (4390844), 5-HTR7 (4392422; s7062), and VASP (4392422; S14748) siRNAs were purchased from Thermo Fisher (Waltham, MA). All siRNA transfections were performed using RNAiMAX (5 µL; Thermo Fisher) and OptiMEM (400 µL; Thermo Fisher). 200,000 cells were trypsinized and seeded in 6-well plates in complete media. After cells adhered (∼1 hr), siRNAs in RNAiMAX/OptiMEM were added to cells in complete media (2 mL) at a final concentration of 50 nM. Cells were incubated with siRNAs for 3 days.

### RT-qPCR

Total RNA was isolated from cells using the PureLink RNA Mini Kit (Thermo Fisher; 12183018A) and was used for reverse transcription using a High-Capacity cDNA Reverse Transcription Kit (Thermo Fisher; 4368814). qPCR was performed using PowerUp SYBR Green Master Mix (Thermo Fisher; A25742) on a real-time PCR detection system (CFX96; Bio-Rad). Relative mRNA levels were calculated using the ΔCt method.

### Transmigration

8 µm (Corning; 3422) transmigration filters were incubated with fibronectin (10 µg/mL) (Thermo Fisher; 33016015) for 12 hr at 4 °C. After drying for 30 min and washing twice with PBS, cells were seeded at a density of 25,000/cm^2^ in starve media with BSA (0.1 %) into the upper well. Medium containing either 10% FBS or 10 µM serotonin as the chemoattractant was added to the lower well. After 12 hr in a tissue culture incubator, transmigrated cells were bonded to the bottom of the filter with 70% ethanol for 10 min, dried for 15 min, washed twice with PBS, and incubated with DAPI (0.1 µg/mL) (Sigma-Aldrich; D9542) for 5 min before the insert was placed on a glass coverslip. Stained nuclei were imaged on a ZOE (Bio-Rad; 1450031) fluorescent imager and counted in Fiji (https://fiji.sc/) using the Analyze Particles plugin.

### Proliferation

100,000 cells were seeded in 6-well tissue culture plates, treated as indicated, and cultured for 5 days. Each day, cells were lifted, resuspended in complete media, diluted 1:1 with Trypan Blue (Sigma; 8154), and loaded into cell counting slides (Bio-Rad; 1450011) for quantification using the Automated Cell Counter (Bio-Rad; 1450102).

### 2D confinement

This protocol has been described in detail elsewhere [17]. Briefly, PDMS (24236-10) was purchased from Electron Microscopy Sciences (Hatfield, PA). 2 mL was cured overnight at 37 °C in each well of a 6-well glass bottom plate (Cellvis; P06-1.5H-N). Using a biopsy punch (World Precision Instruments; 504535), an 8 mm hole was cut, and 3 mL of serum-free media containing 1% BSA was added to each well and incubated overnight at 37 °C. After removing the serum-free media containing 1% BSA, 300 μL of complete media containing trypsinized cells (250,000 to 1 million) and 2 μL of 3.11 μm beads (Bangs Laboratories, Fishers, IN; PS05002) were then pipetted into the round opening. The vacuum created by briefly lifting one side of the hole with a 1 mL pipette tip was used to move cells and beads underneath the PDMS. Finally, 3 mL of complete media was added to each well. Cells recovered for ∼60 min before imaging.

### Microchannel preparation

PDMS (Electron Microscopy Sciences; 24236-10) was prepared using a 1:7 base and curing agent ratio. Uncured PDMS was poured over the wafer mold, placed in a vacuum chamber to remove bubbles, moved to a 37 °C incubator, and left to cure overnight. After curing, small PDMS slabs with microchannels were cut using a scalpel, whereas cell loading ports were cut using a 0.4 cm hole punch (Fisher Scientific; 12-460-409).

For making PDMS-coated cover glass (Fisher Scientific; 12-545-81), 30 µL of uncured PDMS was pipetted into the center of the cover glass, placed in a modified mini-centrifuge, and spun for 30 sec for even spreading. The PDMS-coated cover glass was cured for at least 1 hr on a 95 °C hot plate. To bond the slab and coated cover glass, PDMS surfaces were activated for ∼1 min by plasma treatment (Harrick Plasma; PDC-32G). The apparatus was incubated at 37 °C for at least 1 hr for complete bonding.

### Microchannel coating

Before microchannel coating, surfaces were first activated by plasma treatment. VCAM-1 (R&D Systems; 862-VC) or BSA (VWR; VWRV0332) was used for coating at 1 µg/mL and 1%, respectively, in PBS. Immediately after plasma treatment, VCAM-1 or BSA solution was pumped into microchannels using a modified motorized pipette. To remove any bubbles pumped into microchannels, the apparatus was left to coat in a vacuum chamber for at least 1 hr. Microchannels were then rinsed by repeatedly pumping in new PBS. Finally, microchannels were incubated overnight at 4 °C in complete media before use.

### Microchannel loading

Before cells were loaded into microchannels, complete media was aspirated, and microchannels were placed into an interchangeable cover glass dish (Bioptechs; 190310-35). Freshly trypsinized cells in 300 µL of complete media, stained with 1 µL of fluorescent membrane dye (Thermo Fisher; C10046), were pumped into microchannels using a modified motorized pipette. Once at least 20 cells were observed in microchannels by low magnification brightfield microscopy, microchannels were covered with 2 mL of complete media. Before imaging, a lid was placed on the apparatus to prevent evaporation.

### Public transcriptomic data

Transcriptomic data were analyzed by The Cancer Genome Atlas (https://www.cancer.gov/ccg/research/genome-sequencing/tcga) using the skin cutaneous melanoma (SKCM) dataset. These public data were sourced from the Genomic Data Commons (GDC; https://portal.gdc.cancer.gov/) and accessed through cBioPortal (https://www.cbioportal.org/). For survival analyses, patients were stratified according to their median mRNA level (transcripts per million; TPM) for each enzyme.

### Cell classification in 2D confinement

Cells that displayed directionally persistent migration over at least 4 frames (32 min) were classified as leader mobile (LM). Any cell with a large bleb that remained stable for at least 4 frames (32 min) was considered to have a leader bleb.

### Cell classification in microchannels

Cells that moved at least ½ of their original length over a 5 hr timelapse movie were considered motile. Cells that displayed only blebs were classified as amoeboid, whereas cells that displayed only actin-based protrusions were classified as mesenchymal. Cells with blebs and actin-based protrusions were classified as hybrid.

### Morphometric analysis

The Fiji (https://fiji.sc/) plugin, Analyze_Blebs (https://github.com/karlvosatka/analyze_blebs), was used to measure the largest bleb area, aspect ratio, roundness, circularity, solidity, and Feret’s diameter of cells from timelapse movies [18].

### Cell migration

To perform cell speed and directionality ratio analyses, we used a set of macros, DiPer, developed by Gorelik and colleagues, and the Fiji (https://fiji.sc/) plugin, MTrackJ, created by Erik Meijering for manual tracking [33, 34]. Brightfield imaging confirmed that beads or debris were not obstructing the cell.

### FRET

Ratio images of FRET (CFP excitation/YFP emission) to CFP (CFP excitation/CFP emission) were generated and analyzed in Fiji (https://fiji.sc/).

### Cellular contractility and stiffness assay

The previously described gel sandwich assay was used with minor modifications [6]. 30 mm and 18 mm glass coverslips were activated by aminosilanization (Sigma Aldrich; 281778) for 5 min and then treated for 30 min with 0.5% glutaraldehyde (Electron Microscopy Sciences; 16350) before thoroughly washing with PBS. 1 kPa polyacrylamide gels were made using 2 µL of blue fluorescent beads (ThermoFisher; F8805), 18.8 µL of 40% acrylamide solution (Bio-Rad; 161-0140), and 12.5 µL of bis-acrylamide (Bio-Rad; 161-0142) in 250 µL of PBS. Finally, 2.5 µL of Ammonium Persulfate (APS; 10% in water) and 0.5 µL of Tetramethylethylenediamine (TEMED) was added before spreading 9 µL drops onto treated glass under coverslips. After polymerizing for 1 hr, the coverslip was lifted in PBS, rinsed, and incubated overnight in PBS. Before each assay, the gel attached to the 18 mm coverslip was placed on a 14 mm diameter, 2 cm high PDMS column for applying a slight pressure to the coverslip with its own weight. Then, both gels were incubated for 1 hr in media before seeding cells. After the bottom gel in an interchangeable cover glass dish (Bioptechs; 190310-35) was placed on the microscope stage, the PDMS column with the top gel was placed over the cells seeded on the bottom gels, confining cells between the two gels. After 1 hr of adaptation, the height of cells was determined with beads by measuring the distance between gels, whereas the cell diameter was measured using a far-red plasma membrane dye (ThermoFisher; C10046). Stiffness was defined as the height (*h*) divided by the diameter (*d*). If drugs were used, gels were first incubated with drug in media for 1 hr before obtaining measurements.

### Tissue Array Staining

Melanoma tumor arrays (ME1006) were purchased from Tissue Array (Derwood, MD). Tissues were deparaffinized by baking at 60 °C for 1 hr, followed by xylene washes. Tissues were rehydrated using decreasing concentrations of ethanol (25% increments), from 100% to 25% for 5 min each, before rinsing in distilled water. The antigen retrieval step was performed by immersing slides in Tris-EDTA buffer (pH 9.0) at 95 °C for 15 min, followed by cooling the slide at room temperature for 30 min. Slides were then washed using HEPES buffered saline (HBS) supplemented with 0.1% BSA and 2 mM EDTA. Tissues were permeabilized in supplemented HBS with 0.1% Triton X-100 for 10 min. This was followed by a blocking step using HBS with 1% BSA, 1% fish gelatin (Sigma Aldrich; G7765), and 2 mM EDTA for >2 hr at room temperature. The primary antibody, MAOB (Abcam; ab133270), was then added at a 1:50 dilution in HBS supplemented with 0.1% BSA and 2 mM EDTA overnight at 4 °C. The following day, the slide was washed with HBS for 15 min, repeated 3 times, followed by overnight incubation with an anti-rabbit Alexa Fluor 488 conjugated secondary antibody (1:250) (Thermo Fisher; A-21206). The slides were then washed and rinsed with distilled water. Slides were mounted with EverBright TrueBlack Hardset Mounting Medium containing DAPI (Biotium; 23018-T), cured overnight in the dark at room temperature, and then stored at 4 °C before imaging.

### Microscopy

High-resolution imaging was performed using a DeltaVision (Issaquah, WA) Elite imaging system mounted on an Olympus (Tokyo, Japan) IX71 stand with a motorized stage (XYZ), Ultimate Focus, solid state light source, fast filter wheel for DAPI, CFP, FITC, GFP, YFP, TRITC, mCherry, and Cy5 channels, critical illumination, Olympus UPlanSApo 20X/0.75 NA DIC (air), UPlanSApo 30X/1.05 NA DIC (silicone), PlanApo N 60X/1.42 NA DIC (oil), UPlanSApo 60X/1.30 NA DIC (silicone), and UPlanSApo 100X/1.40 NA DIC (oil) objectives, Photometrics (Tucson, AZ) CoolSNAP HQ2 camera, SoftWoRx (Preston, UK) software with constrained iterative deconvolution, cage incubator, and vibration isolation table.

### Statistics

Sample sizes were determined empirically and based on saturation. For comparing relative changes, statistical significance was determined by a one-sample *t*-test (hypothetical value = 1 or 100%). For comparing two groups, statistical significance was determined using either a Welch’s *t*-test (parametric) or a Mann-Whitney test (non-parametric). For comparisons of three or more groups, statistical significance was determined using either a one-way ANOVA followed by Dunnett’s post-hoc test (parametric) or Kruskal-Wallis test followed by Dunn’s post-hoc correction (non-parametric). Outliers were identified using the ROUT method (Q = 1%). Normality was determined by a D’Agostino & Pearson test. Each statistical test was performed in GraphPad (Boston, MA) Prism. For categorical data, χ^2^ tests were used to determine statistical significance. Significance levels were defined as: * - p ≤ 0.05, ** - p ≤ 0.01, *** - p ≤ 0.001, and **** - p ≤ 0.0001

## MOVIES

**Movie S1.** An A375 cell adopting a leader mobile (LM) phenotype during vertical confinement to ∼3 µm beneath a BSA coated (1%) PDMS ceiling.

**Movie S2.** An A375 cell displaying a hybrid phenotype while confined within a VCAM-1 coated (1 µg/mL) channel.

**Movie S3.** Serotonin-induced PKA activation in an A375 cell. The cell was transiently transfected with Eevee-AKAR and treated with serotonin (10 µM) at 5 min. Warmer pseudocolors indicate elevated levels of PKA activity.

**Movie S4.** Serotonin-induced ROCK activation in an A375 cell. The cell was transiently transfected with Eevee-ROCK and stimulated with serotonin (10 µM) at 5 min. Warmer pseudocolors indicate elevated levels of ROCK activity.

---

**Supplemental Figure 1.**
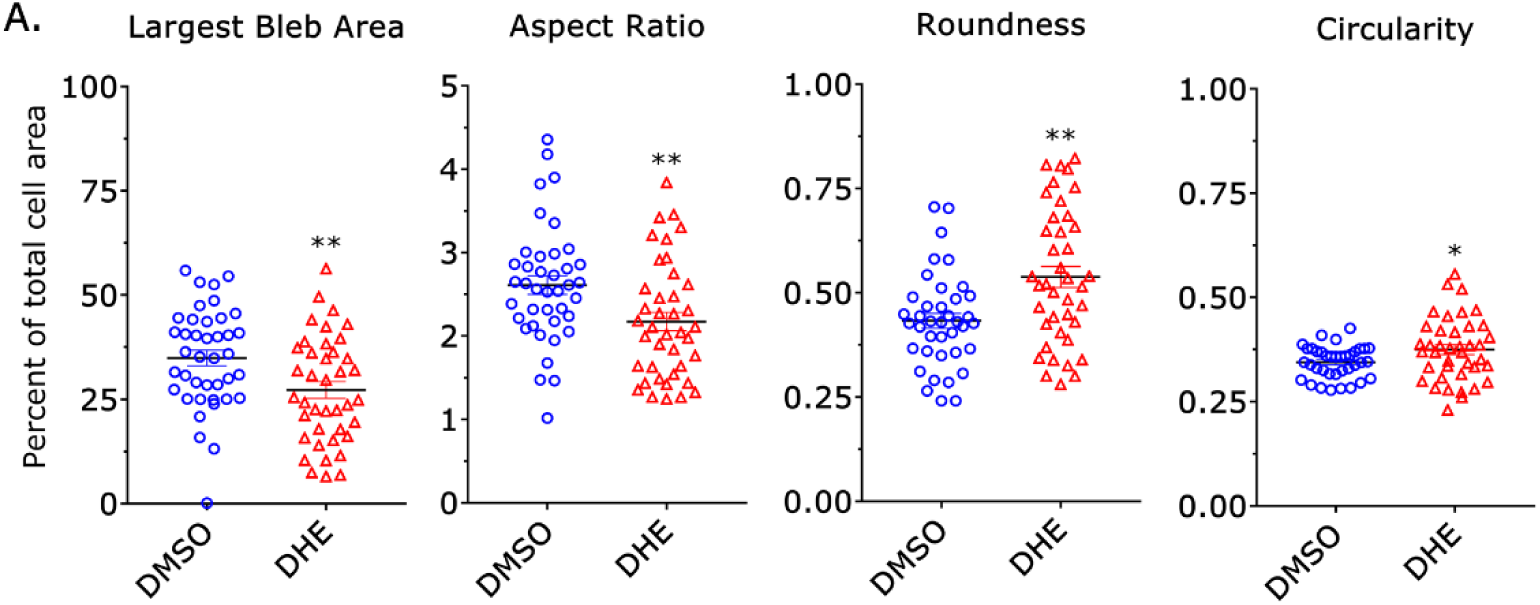
Morphometric profiling of DHE treated cells. **A.** The largest bleb area, aspect ratio, and roundness of A375 cells treated for 1 hr with DMSO or DHE (10 µM). To induce leader bleb-based migration, cells were vertically confined down to ∼3 µm by a BSA coated (1%) PDMS ceiling. Data are presented as the mean +/- SEM. Statistical significance was determined by a Welch’s *t*-test. Significance levels: * - p ≤ 0.05, ** - p ≤ 0.01, *** - p ≤ 0.001, and **** - p ≤ 0.0001.

**Supplemental Figure 2.**
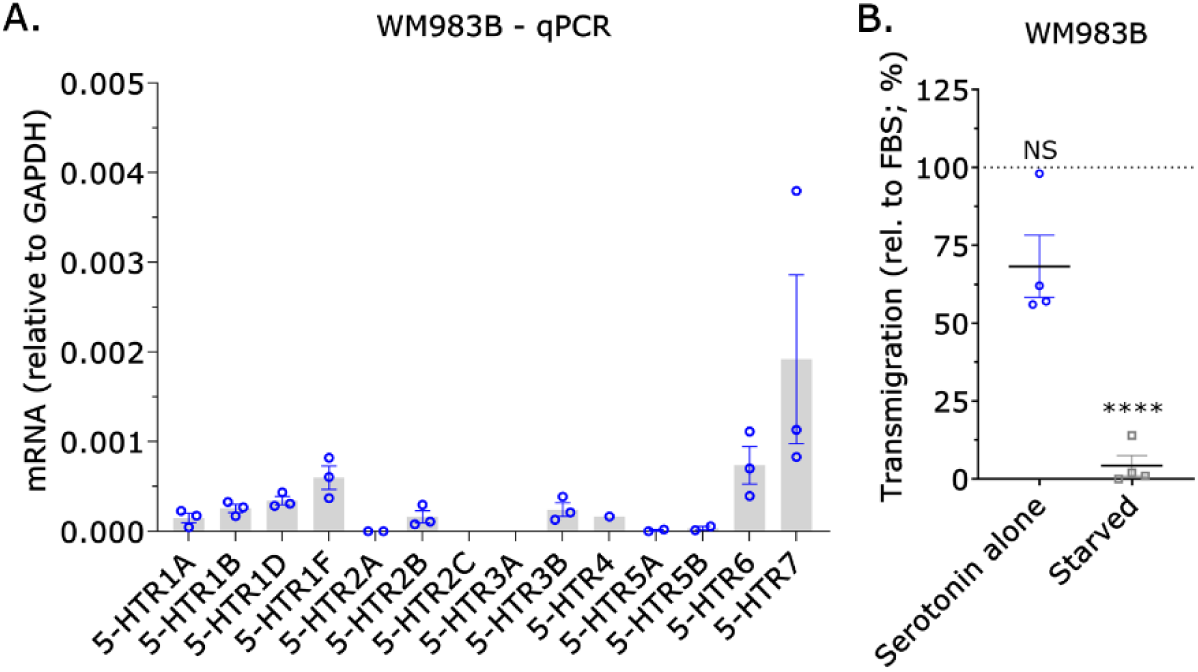
Abundant levels of the 5-HTR7 transcript are found in an additional patient derived melanoma cell line. **A.** The 5-HTR7 transcript is abundant in WM983B cells, as quantified by reverse transcriptase qPCR. Data represent the mean +/- SEM. **B.** Serotonin stimulates cell transmigration through 8 µm pores, relative to serum (10%) and starved controls. Data represent the mean +/- SEM. Statistical significance was determined by a Welch’s *t*-test. Significance levels: * - p ≤ 0.05, ** - p ≤ 0.01, *** - p ≤ 0.001, and **** - p ≤ 0.0001.

**Supplemental Figure 3.**
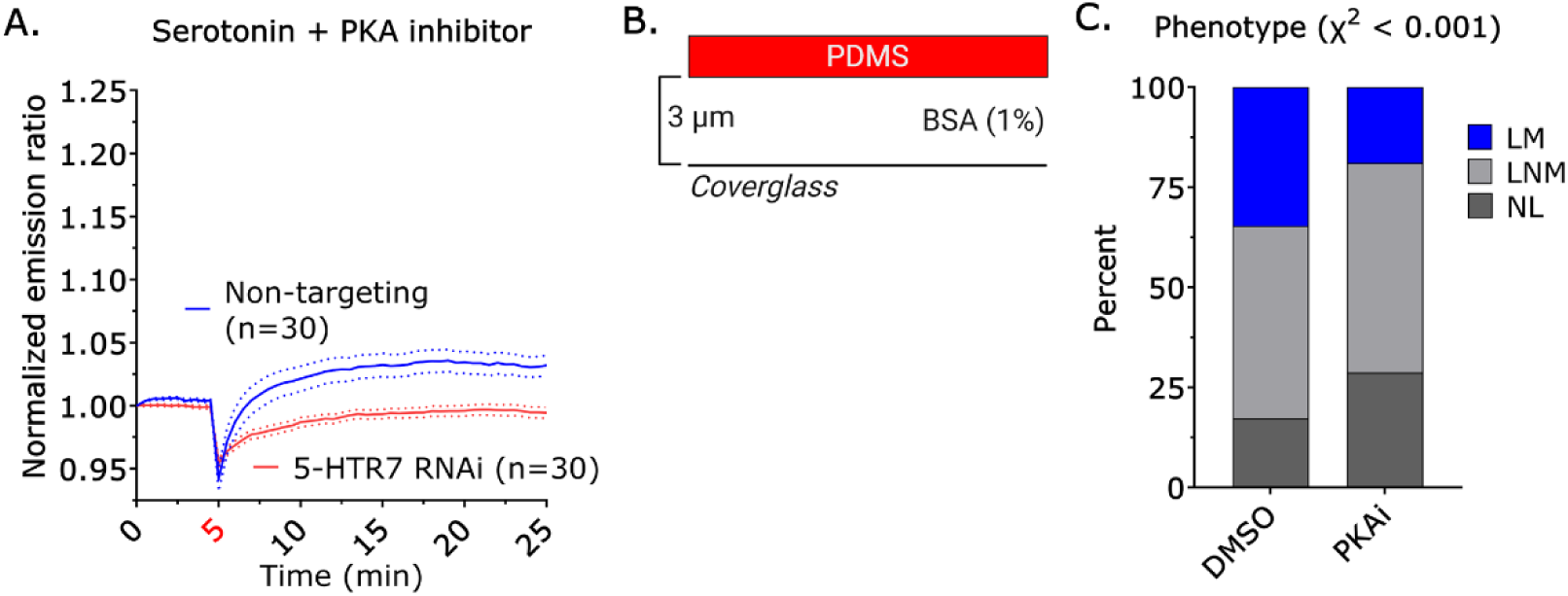
PKA activity drives leader bleb-based migration. **A.** Co-adminstration of a PKA inhibitor (KT5720; 10 µM) and serotonin (10 µM) prevents the induction of kinase activity in A375 cells treated with a non-targeting siRNA, relative to a 5-HTR7 siRNA. Compounds were added to serum-starved cells at 5 min. Data are shown as the mean +/- SEM. **B.** Vertical confinement setup used to induce leader bleb-based migration. Cells are compressed beneath a BSA coated (1%) PDMS ceiling, maintained at a defined height using 3 µm beads. **C.** Distribution of leader mobile (LM), leader non-mobile (LNM), and no leader (NL) phenotypes in cells treated with DMSO (*n* = 426) or PKA inhibitor (KT5720; 10 µM) (*n* = 377). Statistical significance was determined using a χ^2^ test. Significance levels: * - p ≤ 0.05, ** - p ≤ 0.01, *** - p ≤ 0.001, and **** - p ≤ 0.0001.

**Supplemental Figure 4.**
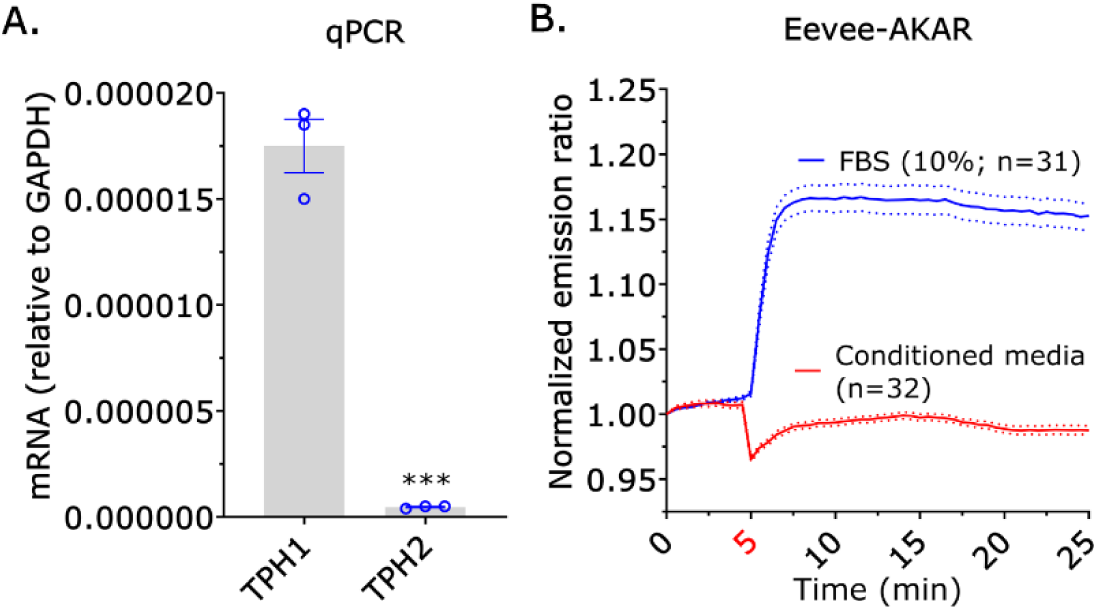
Melanoma cells rely on environmental serotonin rather than autocrine production. **A.** TPH1 and TPH2 transcript levels are minimal in A375 cells, quantified by reverse transcriptase qPCR. Data represent the mean +/- SEM. Statistical differences were determined by a Welch’s *t*-test. **B.** The addition of conditioned media is unable to induce cAMP/PKA signaling, in contrast to the robust response following the addition of serum (10%). Treatments were applied to serum-starved cells at 5 min. Data represent the mean +/- SEM. Significance levels: * - p ≤ 0.05, ** - p ≤ 0.01, *** - p ≤ 0.001, and **** - p ≤ 0.0001.

**Supplemental Figure 5.**
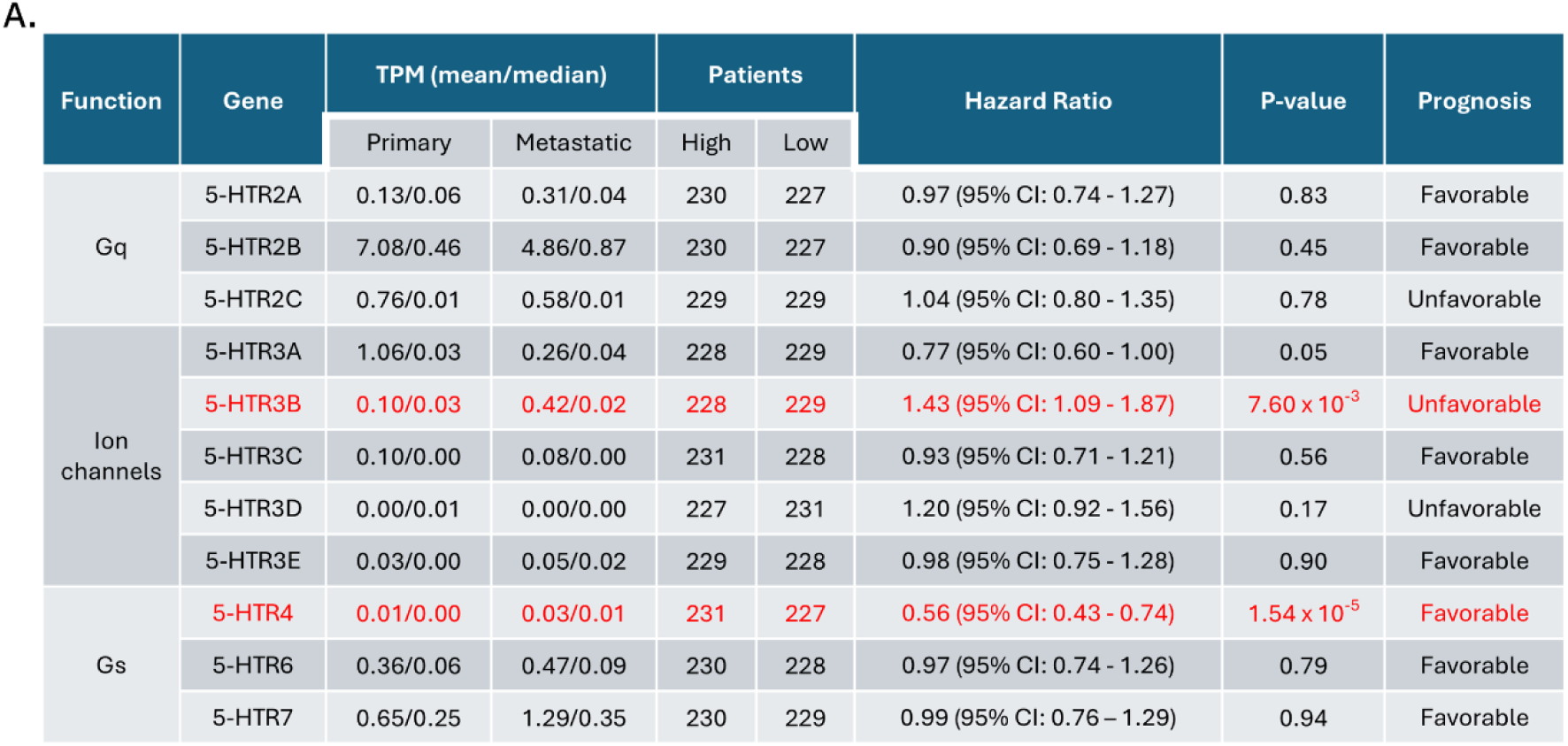
Transcriptomic profiling of serotonin receptors in cutaneous melanoma patients. **A.** Tabulation of Gq coupled, ion channel, and Gs coupled serotonin receptors, detailing transcripts per million (TPM) values within primary and metastatic tumors. The table provides the number of patients stratified by receptor transcript abundance, along with hazard ratios, p-values, and overall disease prognoses. Red highlighting indicates receptors with a significant hazard ratio (p ≤ 0.05). Transcriptomic data for cutaneous melanoma were obtained from The Cancer Genome Atlas through the Genomic Data Commons.

**Supplemental Figure 6.**
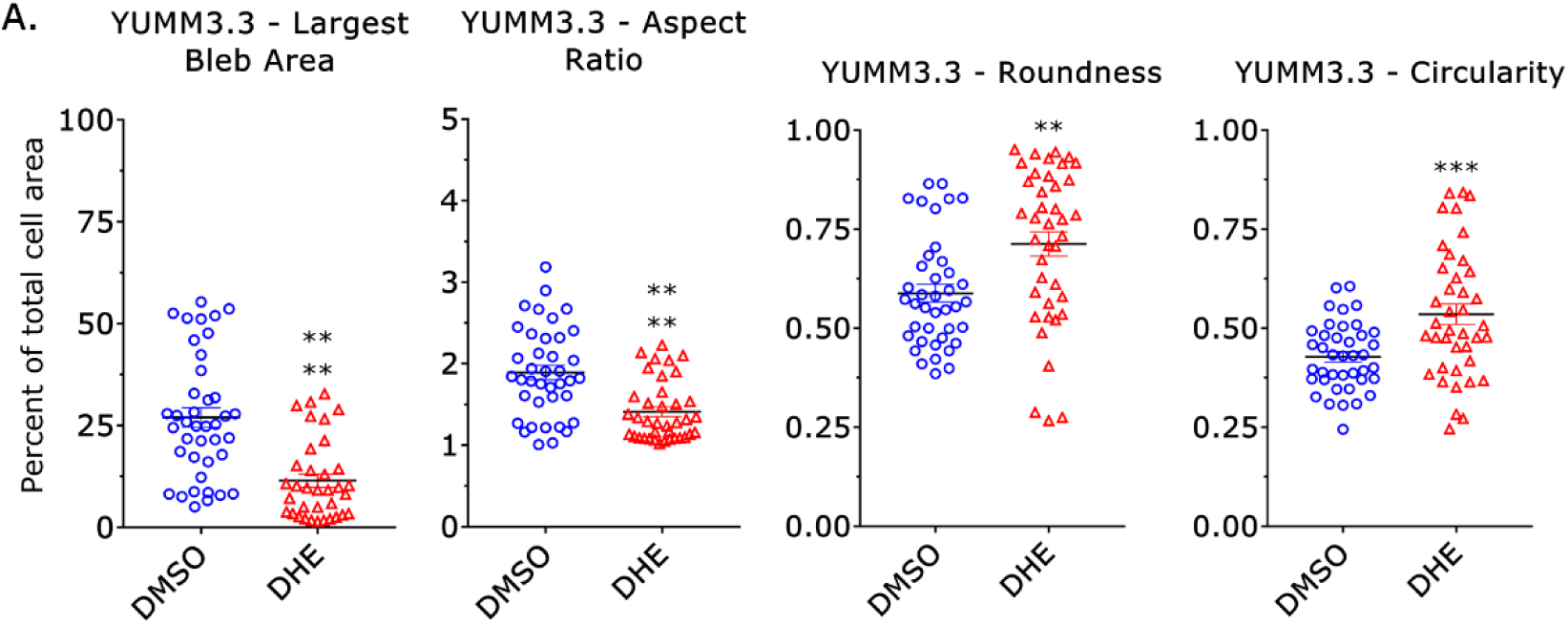
Morphometric profiling of DHE treated YUMM3.3 cells. **A.** Quantification of largest bleb area, aspect ratio, and roundness of cells treated for 1 hr with DMSO or DHE (10 µM). To induce leader bleb-based migration, cells were vertically confined down to ∼3 µm by a BSA coated (1%) PDMS ceiling. Data are presented as the mean +/- SEM. Statistical significance was determined by a Welch’s *t*-test. Significance levels: * - p ≤ 0.05, ** - p ≤ 0.01, *** - p ≤ 0.001, and **** - p ≤ 0.0001.

## Notes

### Competing Interest Statement

The authors have declared no competing interest.

## References

1 Yamada KM, Sixt M. Mechanisms of 3D cell migration. Nat Rev Mol Cell Biol 2019; 20: 738–752.

2 George S, Martin JAJ, Graziani V, Sanz-Moreno V. Amoeboid migration in health and disease: Immune responses versus cancer dissemination. Front Cell Dev Biol 2022; 10: 1091801.

3 Charras G, Paluch E. Blebs lead the way: how to migrate without lamellipodia. Nat Rev Mol Cell Biol 2008; 9: 730–736.

4 Driscoll MK, Welf ES, Weems A, Sapoznik E, Zhou F, Murali VS et al. Proteolysis-free amoeboid migration of melanoma cells through crowded environments via bleb-driven worrying. Dev Cell 2024; 59: 2414–2428 e2418.

5 Logue JS, Cartagena-Rivera AX, Baird MA, Davidson MW, Chadwick RS, Waterman CM. Erk regulation of actin capping and bundling by Eps8 promotes cortex tension and leader bleb-based migration. eLife 2015; 4.

6 Liu YJ, Le Berre M, Lautenschlaeger F, Maiuri P, Callan-Jones A, Heuze M et al. Confinement and low adhesion induce fast amoeboid migration of slow mesenchymal cells. Cell 2015; 160: 659–672.

7 Ruprecht V, Wieser S, Callan-Jones A, Smutny M, Morita H, Sako K et al. Cortical contractility triggers a stochastic switch to fast amoeboid cell motility. Cell 2015; 160: 673–685.

8 Bergert M, Erzberger A, Desai RA, Aspalter IM, Oates AC, Charras G et al. Force transmission during adhesion-independent migration. Nat Cell Biol 2015; 17: 524–529.

9 Bergert M, Chandradoss SD, Desai RA, Paluch E. Cell mechanics control rapid transitions between blebs and lamellipodia during migration. Proc Natl Acad Sci U S A 2012; 109: 14434–14439.

10 Tozluoglu M, Tournier AL, Jenkins RP, Hooper S, Bates PA, Sahai E. Matrix geometry determines optimal cancer cell migration strategy and modulates response to interventions. Nat Cell Biol 2013; 15: 751–762.

11 Cantelli G, Orgaz JL, Rodriguez-Hernandez I, Karagiannis P, Maiques O, Matias-Guiu X et al. TGF-β-induced transcription sustains amoeboid melanoma migration and dissemination. Curr Biol 2015; 25: 2899–2914.

12 Kar N, Caruso AP, Prokopiou N, Abrenica A, Logue JS. The activation of INF2 by Piezo1/Ca^2+^ is required for mesenchymal-to-amoeboid transition in confined environments. Curr Biol 2025; 35, 1791–1804.e5.

13 Wales P, Schuberth CE, Aufschnaiter R, Fels J, Garcia-Aguilar I, Janning A et al. Calcium-mediated actin reset (CaAR) mediates acute cell adaptations. eLife 2016; 5.

14 Logue JS, Cartagena-Rivera AX, Chadwick RS. c-Src activity is differentially required by cancer cell motility modes. Oncogene 2018; 37: 2104–2121.

15 Ullo MF, Logue JS. Re-thinking preclinical models of cancer metastasis. Oncoscience 2018; 5: 252–253.

16 Kuang S, Abrenica A, Kar N, Logue JS. Targeting Cholesterol-Dependent Piezo1 Activation Impairs Amoeboid Migration in Melanoma Cells. bioRxiv 2025.

17 Logue JS, Waterman CM, Chadwick RS. A simple method for precisely controlling the confinement of cells in culture. Protocol Exchange 2018; doi:10.1038/protex.2018.033.

18 Vosatka KW, Lavenus SB, Logue JS. A novel Fiji/ImageJ plugin for the rapid analysis of blebbing cells. PLoS One 2022; 17: e0267740.

19 Kroeze WK, Sassano MF, Huang XP, Lansu K, McCorvy JD, Giguere PM et al. PRESTO-Tango as an open-source resource for interrogation of the druggable human GPCRome. Nat Struct Mol Biol 2015; 22: 362–369.

20 Lovenberg TW, Baron BM, de Lecea L, Miller JD, Prosser RA, Rea MA et al. A novel adenylyl cyclase-activating serotonin receptor (5-HT7) implicated in the regulation of mammalian circadian rhythms. Neuron 1993; 11: 449–458.

21 Kvachnina E, Liu G, Dityatev A, Renner U, Dumuis A, Richter DW et al. 5-HT7 receptor is coupled to G alpha subunits of heterotrimeric G12-protein to regulate gene transcription and neuronal morphology. J Neurosci 2005; 25: 7821–7830.

22 Komatsu N, Aoki K, Yamada M, Yukinaga H, Fujita Y, Kamioka Y et al. Development of an optimized backbone of FRET biosensors for kinases and GTPases. Mol Biol Cell 2011; 22: 4647–4656.

23 Walther DJ, Peter JU, Bashammakh S, Hortnagl H, Voits M, Fink H et al. Synthesis of serotonin by a second tryptophan hydroxylase isoform. Science 2003; 299: 76.

24 Curtis JJ, Seymour CB, Mothersill CE. Cell Line-Specific Direct Irradiation and Bystander Responses are Influenced by Fetal Bovine Serum Serotonin Concentrations. Radiat Res 2018; 190: 262–270.

25 Li C, Imanishi A, Komatsu N, Terai K, Amano M, Kaibuchi K et al. A FRET Biosensor for ROCK Based on a Consensus Substrate Sequence Identified by KISS Technology. Cell Struct Funct 2017; 42: 1–13.

26 Butt E, Abel K, Krieger M, Palm D, Hoppe V, Hoppe J et al. cAMP- and cGMP-dependent protein kinase phosphorylation sites of the focal adhesion vasodilator-stimulated phosphoprotein (VASP) in vitro and in intact human platelets. J Biol Chem 1994; 269: 14509–14517.

27 Meeth K, Wang JX, Micevic G, Damsky W, Bosenberg MW. The YUMM lines: a series of congenic mouse melanoma cell lines with defined genetic alterations. Pigment Cell Melanoma Res 2016; 29: 590–597.

28 Jin H, Oksenberg D, Ashkenazi A, Peroutka SJ, Duncan AM, Rozmahel R et al. Characterization of the human 5-hydroxytryptamine1B receptor. J Biol Chem 1992; 267: 5735–5738.

29 Wang C, Jiang Y, Ma J, Wu H, Wacker D, Katritch V et al. Structural basis for molecular recognition at serotonin receptors. Science 2013; 340: 610–614.

30 Mawe GM, Hoffman JM. Serotonin signalling in the gut--functions, dysfunctions and therapeutic targets. Nat Rev Gastroenterol Hepatol 2013; 10: 473–486.

31 Chen L, Huang S, Wu X, He W, Song M. Serotonin signalling in cancer: Emerging mechanisms and therapeutic opportunities. Clin Transl Med 2024; 14: e1750.

32 Barzik M, Kotova TI, Higgs HN, Hazelwood L, Hanein D, Gertler FB et al. Ena/VASP proteins enhance actin polymerization in the presence of barbed end capping proteins. J Biol Chem 2005; 280: 28653–28662.

33 Meijering E, Dzyubachyk O, Smal I. Methods for cell and particle tracking. Methods Enzymol 2012; 504: 183–200.

34 Gorelik R, Gautreau A. Quantitative and unbiased analysis of directional persistence in cell migration. Nat Protoc 2014; 9: 1931–1943.

